# Susceptibility of swine cells and domestic pigs to SARS-CoV-2

**DOI:** 10.1101/2020.08.15.252395

**Authors:** David A. Meekins, Igor Morozov, Jessie D. Trujillo, Natasha N. Gaudreault, Dashzeveg Bold, Bianca L. Artiaga, Sabarish V. Indran, Taeyong Kwon, Velmurugan Balaraman, Daniel W. Madden, Heinz Feldmann, Jamie Henningson, Wenjun Ma, Udeni B. R. Balasuriya, Juergen A. Richt

## Abstract

The emergence of SARS-CoV-2 has resulted in an ongoing global pandemic with significant morbidity, mortality, and economic consequences. The susceptibility of different animal species to SARS-CoV-2 is of concern due to the potential for interspecies transmission, and the requirement for pre-clinical animal models to develop effective countermeasures. In the current study, we determined the ability of SARS-CoV-2 to (i) replicate in porcine cell lines, (ii) establish infection in domestic pigs via experimental oral/intranasal/intratracheal inoculation, and (iii) transmit to co-housed naive sentinel pigs. SARS-CoV-2 was able to replicate in two different porcine cell lines with cytopathic effects. Interestingly, none of the SARS-CoV-2-inoculated pigs showed evidence of clinical signs, viral replication or SARS-CoV-2-specific antibody responses. Moreover, none of the sentinel pigs displayed markers of SARS-CoV-2 infection. These data indicate that although different porcine cell lines are permissive to SARS-CoV-2, five-week old pigs are not susceptible to infection via oral/intranasal/intratracheal challenge. Pigs are therefore unlikely to be significant carriers of SARS-CoV-2 and are not a suitable pre-clinical animal model to study SARS-CoV-2 pathogenesis or efficacy of respective vaccines or therapeutics.

## Introduction

The emergence of SARS-CoV-2, the causative agent of COVID-19, has resulted in a global pandemic with over 20 million cases and 740,000 deaths as of August 13, 2020 [1,2]. SARS-CoV-2 causes a respiratory disease in humans with a broad clinical presentation, ranging from asymptomatic or mild illness to severe fatal disease with multi-organ failure [3-6]. SARS-CoV-2 is rapidly transmissible via contact with infected respiratory droplets and can also be transmitted by asymptomatic carriers [6-8]. To curb viral spread, countries have instituted varying levels of social distancing policies, which have significant negative economic and social impacts [9]. Mitigating the effects of this unprecedented pandemic will necessitate the development of effective vaccines and therapeutics, which will require well-characterized and standardized pre-clinical animal models.

SARS-CoV-2 is a member of the *Betacoronavirus* genus that includes the pathogenic human viruses SARS-CoV-1 and MERS-CoV [2,10-12]. While details of the origin of SARS-CoV-2 are unknown, evidence indicates it emerged from a zoonotic spillover event, with bats and perhaps pangolins as probable origin species [2,13-15]. The potential for a reverse zoonotic event, i.e. human-to-animal transmission, is possible and of significant concern to animal and public health [16-18]. Instances of natural human-to-animal transmission of SARS-CoV-2 have been reported with COVID-19 patients in domestic settings (dogs and cats), zoos (lions and tigers), and farms (mink) [18-20]. Therefore, investigations into the infectivity of SARS-CoV-2 in various animal species with human contact are essential to assess and control the risk of a spillover event and to establish the role these animals may play in the ecology of the virus.

Several studies have determined the susceptibility of different animal species to SARS-CoV-2 via experimental infection [20,21]. Cats, hamsters, and ferrets are highly susceptible to SARS-CoV-2 infection, demonstrate varying clinical and pathological disease manifestations, readily transmit the virus to naïve animals, and mount a virus-specific immune response [22-28]. Dogs are mildly susceptible to experimental SARS-CoV-2 infection, with limited viral replication but with clear evidence of seroconversion in some animals [22]. Poultry species seem to be resistant to SARS-CoV-2 infection [22,26]. These findings establish the respective utility of different animal species as pre-clinical models to study SARS-CoV-2.

Several lines of evidence suggest that pigs could be susceptible to SARS-CoV-2 infection. Pigs are susceptible to both experimental and natural infection with the related Betacoronavirus, SARS-CoV-1, and demonstrate seroconversion [29,30]. Structure-based analyses predict that the SARS-CoV-2 Spike (S) protein receptor binding domain (RBD) binds the pig angiotensin-converting enzyme 2 (ACE2) entry receptor with similar efficiency compared to human ACE2 [31]. Single-cell screening also indicated that pigs co-express ACE2 and the TMPRSS2 activating factor in a variety of different cell lines, and SARS-CoV-2 replicates in various pig cell lines [2,26,32,33]. Despite these preliminary data indicating that pigs could be susceptible to SARS-CoV-2 infection, two recent studies revealed that intranasal inoculation of three and twelve pigs, respectively, with 10^5^ pfu or TCID_50_ of SARS-CoV-2 did not lead to any detectable viral replication or seroconversion [22,26]. However, the single route of intranasal inoculation used in these studies suggests that additional investigations are necessary before definitive conclusions can be made regarding susceptibility of pigs to SARS-CoV-2.

In the present study, we determined the susceptibility of swine cell lines and domestic pigs to SARS-CoV-2 infection. Two different porcine cell lines were found to be permissive to SARS-CoV-2 infection showing cytopathic effects (CPE). Domestic pigs were challenged via simultaneous oral/intranasal/intratracheal inoculation with a 10^6^ TCID_50_ dose of SARS-CoV-2. SARS-CoV-2 did not replicate in pigs and none of them seroconverted. Furthermore, the virus was not transmitted from SARS-CoV-2 inoculated animals to sentinels. The present findings, combined with the other studies [22,26], confirm that pigs seem resistant to SARS-CoV-2 infection despite clear susceptibility of porcine cell lines. Pigs are therefore unlikely to play an important role in the COVID-19 pandemic as a virus reservoir or as a pre-clinical animal model to study SARS-CoV-2 pathogenesis or develop novel countermeasures.

## Materials and Methods

### Virus and cells

SARS-CoV-2 USA-WA1/2020 isolate (GenBank accession # MN985325) [34] was obtained from BEI resources (catalog # NR-52281, American Type Culture Collection [ATCC^®^]. Manassas, VA, USA). The virus was passaged three times in VeroE6 cells (ATCC^®^ CRL-1586™), before being passaged two times in swine testicle (ST; ATCC CRL-1746™) and four times in porcine kidney (PK-15; ATCC^®^ CCL-33™) cell lines to investigate suitability of these swine cell lines for propagation of SARS-CoV-2. The first passage of the SARS-CoV-2 virus from ST cells was used to prepare the challenge material. Back-titer of the virus inoculum stock was 3.15×10^5^ TCID_50_/mL All experiments involving the SARS-CoV-2 virus were performed under Biosafety Level 3+ (BSL-3+) conditions at the Biosecurity Research Institute (BRI) at Kansas State University (KSU), Manhattan, KS, USA.

### Outline of animal experiments

Animal infection experiments using swine were performed under BSL-3Ag conditions at the BRI at KSU. Animal research was conducted in compliance with the Animal Welfare Act and other federal statutes and regulations relating to animal care and experimentation under protocol #4390, approved by the Institutional Animal Care and Use Committee (IACUC) at Kansas State University on April 8, 2020.

Eighteen pigs (mix of males and females, five weeks of age) were used in the study. Pigs were acquired from a source guaranteed free of swine influenza virus (SIV), porcine circovirus-2 (PCV-2), and porcine reproductive and respiratory syndrome virus (PRRSV) infection. The study outline is illustrated in Figure 1. Upon arrival, pigs were acclimated for 3 days prior to SARS-CoV-2 inoculation. Nine pigs were designated as uninfected negative controls and housed in separate BSL-2 facilities. Three of these uninfected negative control pigs were humanely euthanized at 3 days post challenge (DPC) to provide negative control clinical and tissue samples. The nine principal infected pigs were housed in the same room in two separate groups (4 or 5 pigs each; Figure 1) and were infected with 1.3 x10^6^ TCID_50_ of SARS-CoV-2 orally (1 mL), intranasally (1 ml; 0.5 ml each nostril) and intratracheally (2 mL), after being anesthetized with a mixture of telazol/xylazine. Three sentinel contact pigs were added to each group on 1 DPC (Figure 1). Rectal temperature and signs of clinical disease for each pig were determined daily throughout the study; clinical signs include overall activity/attitude (signs of depression, decreased alertness or unresponsiveness), appetite (based on interest in treats), respiratory signs (sneezing, coughing, labored breathing, nasal discharge), and digestive signs (diarrhea or vomiting). Blood samples and nasal, oropharyngeal, and rectal swabs were collected in virus transport medium (VTM; DMEM plus antibiotic/antimycotic) at 0, 1, 3, 5, 7, 10, 14, and 21 DPC. As summarized in Table 1 and Figure 1, pigs were humanely euthanized for scheduled *post mortem* examinations on 4 DPC (3 principal infected pigs), 8 DPC (3 principal infected pigs), and 21 DPC (remaining 3 principal infected and 6 sentinel pigs) to collect respiratory tissue samples, with blood and swab samples collected on pigs prior to euthanasia. Gross pathological examinations on major organs were performed and respiratory tissue samples were collected and either stored in 10% neutral-buffered formalin or stored as fresh samples at −80°C. Blood and swab samples were all filtered using a 0.2μm filter prior to storage at −80°C.

**Table 1.** Animal groups.

| Group | Treatment | Pig ID#s | Necropsy |
| --- | --- | --- | --- |
| 1 | Principal | 807, 161, 168 | 4 DPC |
| 1 | Principal | 803, 841, 211 | 8 DPC |
| 1 | Principal | 893, 193, 219 | 21 DPC |
| 2 | Sentinel | 848, 194, 222, 851, 172, 894 | 21 DPC |
| 3 | Mock | 201, 183, 238 | 3 DPC (not infected) |

**Figure 1.**
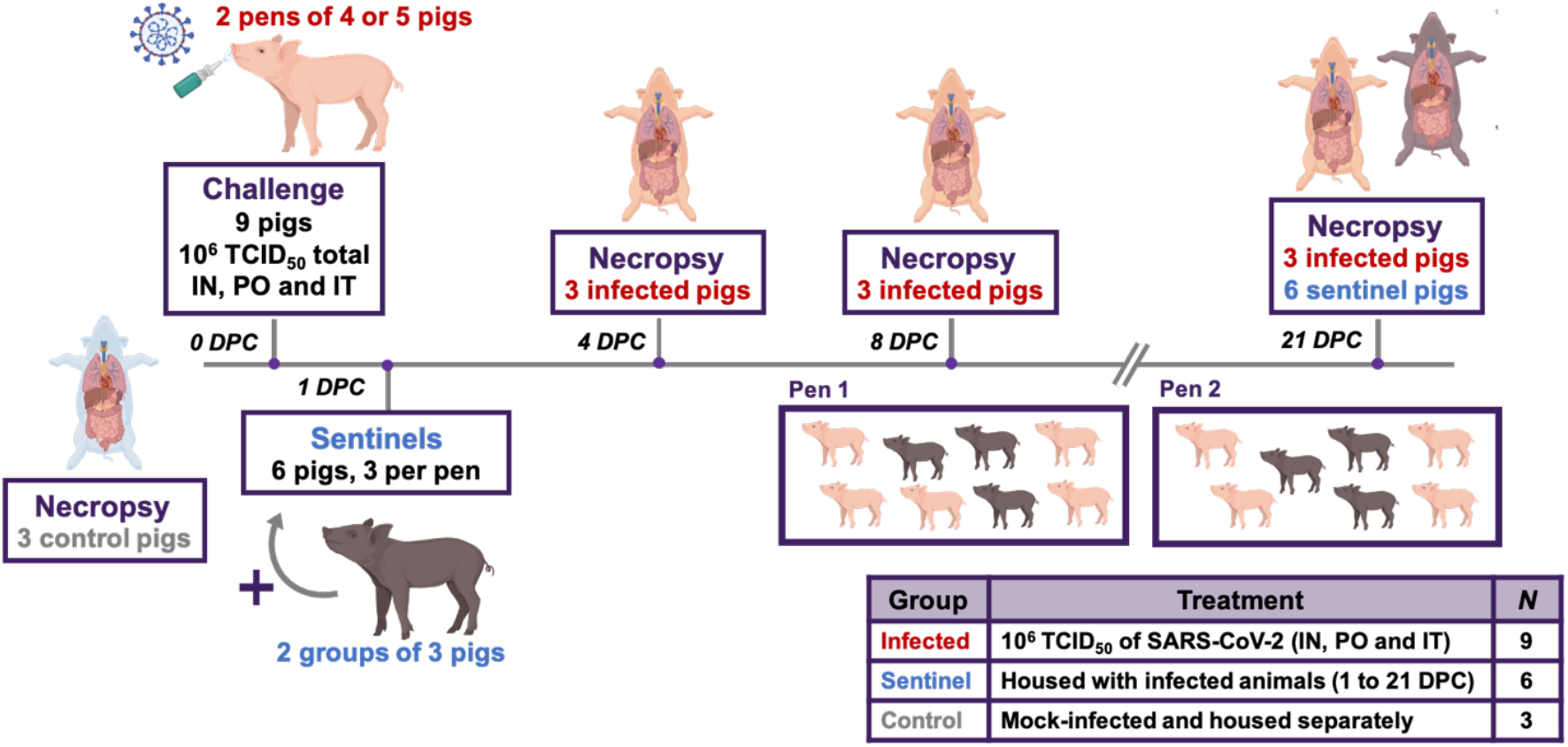
Study Design. Eighteen pigs were placed into three groups. Group 1 (principal infected animals) consisted of nine pigs (four and five in each pen) and was inoculated via intranasal (IN), oral (PO), and intratracheal (IT) routes simultaneously with a total dose of 1 x 10^6^ TCID_50_ of SARS-CoV-2 in 4mL DMEM. The pigs in Group 2 (n=6; sentinel contact animals) and Group 3 (n=3; mock control animals) were housed in a separate room. At 1-day post challenge (DPC), the six pigs in Group 2 were co-mingled with the principal infected animals in Group 1 (three pigs per pen) and served as sentinel contact controls. The remaining three pigs in Group 3 remained in separate housing and served as mock-infected negative controls and were euthanized and necropsied on 3 DPC. Principal infected animals were euthanized and necropsied at 4 (n=3), 8 (n=3), and 21 (n=3) to determine the course of infection. All six sentinel pigs were also euthanized on 21 DPC.

### RNA extraction and reverse transcription quantitative PCR (RT-qPCR)

RNA was isolated from blood, swabs, and tissue samples, using a magnetic bead-based protocol in a BSL-3+ laboratory at the BRI at KSU. Lung tissue homogenates (200 mg per 1mL DMEM; 20% w/v) were prepared by thawing tissue, mincing it into 1mm sections, followed by lysis in a 2 mL sure-lock tube containing 5 mm stainless steel homogenization beads using the TissueLyser LT (Qiagen, Germantown, MD, USA) for 30 seconds at 30 hz followed by 1 min of 30 hz while keeping the sample cold. Following clarification via a 3-minute centrifugation (3,000xg; room temperature), supernatants were mixed with an equal volume of RLT lysis buffer. Blood and clinical swabs were directly mixed with an equal volume of RLT lysis buffer. 200 μL of each sample lysate was used to extract RNA using a magnetic bead-based nucleic acid extraction kit (GeneReach USA, Lexington, MA) on an automated TacoMini™ nucleic acid extraction system (GeneReach USA, Lexington, MA) according to manufacturer’s protocol with the following modifications: beads were added to the sample well, followed by the RLT sample lysate, then 200 μL molecular grade isopropanol (ThermoFisher Scientific, Waltham, MA, USA) was added. The last wash buffer was replaced by molecular grade 200 proof ethanol (ThermoFisher Scientific, Waltham, MA, USA). Extraction positive controls (IDT, IA, USA; 2019-nCoV_N_Positive Control, diluted 1:100 in RLT buffer) and negative controls were employed. Extracted RNA was eluted in 100μL buffer.

Reverse transcription quantitative PCR (RT-qPCR) was performed to detect viral RNA using the CDC standard N2-based SARS-CoV-2 detection assay [35] that was validated for use with the qScript XLT 1-Step RT-qPCR ToughMix (Quantabio, Beverly, MA, USA) on a CFX96 real-time thermocycler (Bio-Rad, Hercules, CA, USA) using a 20 minute RT step and 45 cycle qPCR in a 20 μL reaction volume. RT-qPCR on each sample was performed in duplicate wells with a quantitated PCR positive control (IDT 2019-nCoV N Positive Control, diluted 1:100) and four non-template negative controls on every plate. A positive Ct cut-off of 40 cycles was used. A plasmid template including the SARS-CoV-2 N gene was used as a positive PCR amplification control. A 10-point standard curve using quantitated stock viral RNA (USA-WA1/2020 isolate) was used to quantify RNA copy number.

### Gross pathology and histopathology

During *post mortem* examinations, the upper and lower respiratory tract, central nervous system, lymphatic and cardiovascular systems, gastrointestinal and urogenital systems, and integument were evaluated. Lungs were removed *in toto* and the percentage of the lung surface that was affected by macroscopic lesions was estimated by single veterinarian experienced in evaluating gross porcine lung pathology as previously described [36,37]. Lungs were evaluated for gross pathology such as edema, congestion, discoloration, atelectasis, and consolidation. Tissue samples of interest were collected and either fixed in 10% neutral-buffered formalin for histopathological examination or frozen at −80°C for RT-qPCR testing. Tissues were fixed in formalin for 7 days, then transferred to 70% ethanol (ThermoFisher Scientific, Waltham, MA, USA) prior to trimming and paraffin embedding following standard automated protocols used in the histology section of the Kansas State Veterinary Diagnostic Laboratory. Following embedding, tissue sections were cut and stained with hematoxylin and eosin and evaluated by a board-certified veterinary pathologist who was blinded to the treatment groups.

### Serological testing

To detect SARS-CoV-2 antibodies in sera, indirect ELISAs were performed using recombinant SARS-CoV-2 Receptor Binding Domain (RBD) expressed in HEK cells with a C-terminal Strep-Tag and the Nucleocapsid (N) protein expressed in *E.coli* with a C-terminal His-tag. Briefly, the recombinant proteins were produced in their respective expression systems according to standard procedures and purified using either Ni-NTA (ThermoFisher Scientific, Waltham, MA, USA) or Strep-Tactin (IBA Lifesciences, Goettingen, Germany) columns according to manufacturer’s instructions. For testing, 96-well plates were coated with 100 ng of the recombinant protein in 100μL of sodium carbonate/bicarbonate buffer and incubated overnight at 4°C. The wells were then washed and blocked with casein blocking buffer. Sera dilutions were diluted 1:200 in 100μL of casein blocking buffer, incubated at room temperature for 1 hour, then washed with washing solution. 100 μL anti-pig-IgG or IgM secondary antibodies, conjugated with peroxidase, were diluted 1:2,500 in blocking buffer and then incubated at room temperature for 1 hour, protected from light. TMB colorimetric substrate was added and incubated at room temperature for 5 minutes, then the reaction was stopped with a solution of 0.2 sulfuric acid. The optical density (OD) value was measured at 450 nm within 5 minutes of adding the stop solution to quantify the amount of antigen-binding antibody present in the sample. Sera from mock-infected pigs were used as negative controls. Sera collected from SARS-CoV-2 infected cats, from a different study [38], were used as a positive control. Serum from pigs infected with African Swine Fever Virus (ASFV) and the baculovirus-expressed ASFV-p54 antigen were used as a positive control for anti-pig-IgG or IgM secondary antibodies. The cutoff for a sample being called positive was defined by the average OD 0 DPC serum +3x standard deviation.

The presence of virus-neutralizing antibodies in sera was determined via microneutralization assay. Serum samples were diluted 1:10 and heat-inactivated at 56°C for 30 minutes while shaking. Subsequently, 100 μL per well of serum samples in duplicate were subjected to 2-fold serial dilutions starting at 1:20 through 1:2560 in 100μL culture media. 100 μL of 100 TCID_50_ of SARS-CoV-2 was then added to 100 μL of the sera dilutions and incubated for 1 hour at 37°C, followed by culture of the mixture on VeroE6 cells in 96-well plates. Results of the virus neutralization were determined by the appearance of CPE, which was observed under a microscope at 96 hours post inoculation. The neutralizing antibody titer is determined as the reciprocal of the average serum dilution at which no CPE breakthrough in any of the testing wells is observed. Neutralizing sera from SARS-CoV-2-infected cats from a separate study [38] was used as a positive control.

### Next Generation Sequencing

To determine the consensus sequence of the USA-WA/1/2020 virus and to analyze if there were any nucleic acid substitutions in the SARS-CoV-2 virus after passage in porcine cell lines, RNA was extracted from cell culture supernatant as described above. The RNA was then subjected to RT-PCR amplification using a tiled-primer approach to amplify the entire SARS-CoV-2 genome as described previously [39]. Briefly, the PCR amplicons were pooled and subjected to library preparation for Next Generation Sequencing using the Nextera XT library prep kit (Illumina, San Diego, CA, USA). The library was normalized and sequenced using a MiSeq nano v2 2×250 sequencing kit. The sequence was then analyzed by mapping reads to the parent sequence (Genbank accession # MN985325) [34] to generate a consensus sequence.

## Results

### Propagation of SARS-CoV-2 in swine cells

The SARS-CoV-2 USA-WA1/2020 isolate, which was isolated from a human patient in Washington State, USA, was used as the parent stock for the study [34]. The virus stock was passaged 3 times at a multiplicity of infection (MOI) between 0.001 and 0.01 in VeroE6 cells and NGS sequenced before being inoculated onto porcine cells. The consensus sequence of the VeroE6-passaged virus was 100% identical to the GenBank reference sequence (GenBank accession # MN985325). To determine the susceptibility of porcine cell lines to SARS-CoV-2 infection, swine testicle (ST) and porcine kidney (PK-15) cell lines were inoculated with approximately 0.05 MOI of passage 3 of the VeroE6-passaged SARS-CoV-2 USA-WA/1/2020 isolate. No obvious cytopathic effect (CPE) was observed in either cell line during the first passage, however clear CPE was observed in passage two in ST cells and passage four in PK-15 cells (Figure 2). Cell culture supernatant from SARS-CoV-2 passage one and two on ST cells, and passage three on PK-15 cells was collected, and RNA extracted for sequencing. Next generation sequencing was performed to generate a consensus genomic sequence for the ST- and PK-15-passaged SARS-CoV-2 virus. No nucleotide mutations or amino acid substitutions were observed upon passage in the two porcine cell lines. These results indicate that SARS-CoV-2 is able to readily infect porcine kidney and testicle cells without the requirement of obvious genetic adaptation. SARS-CoV-2 from passage 1 in ST cells was used as challenge material for the pig inoculation.

**Figure 2.**
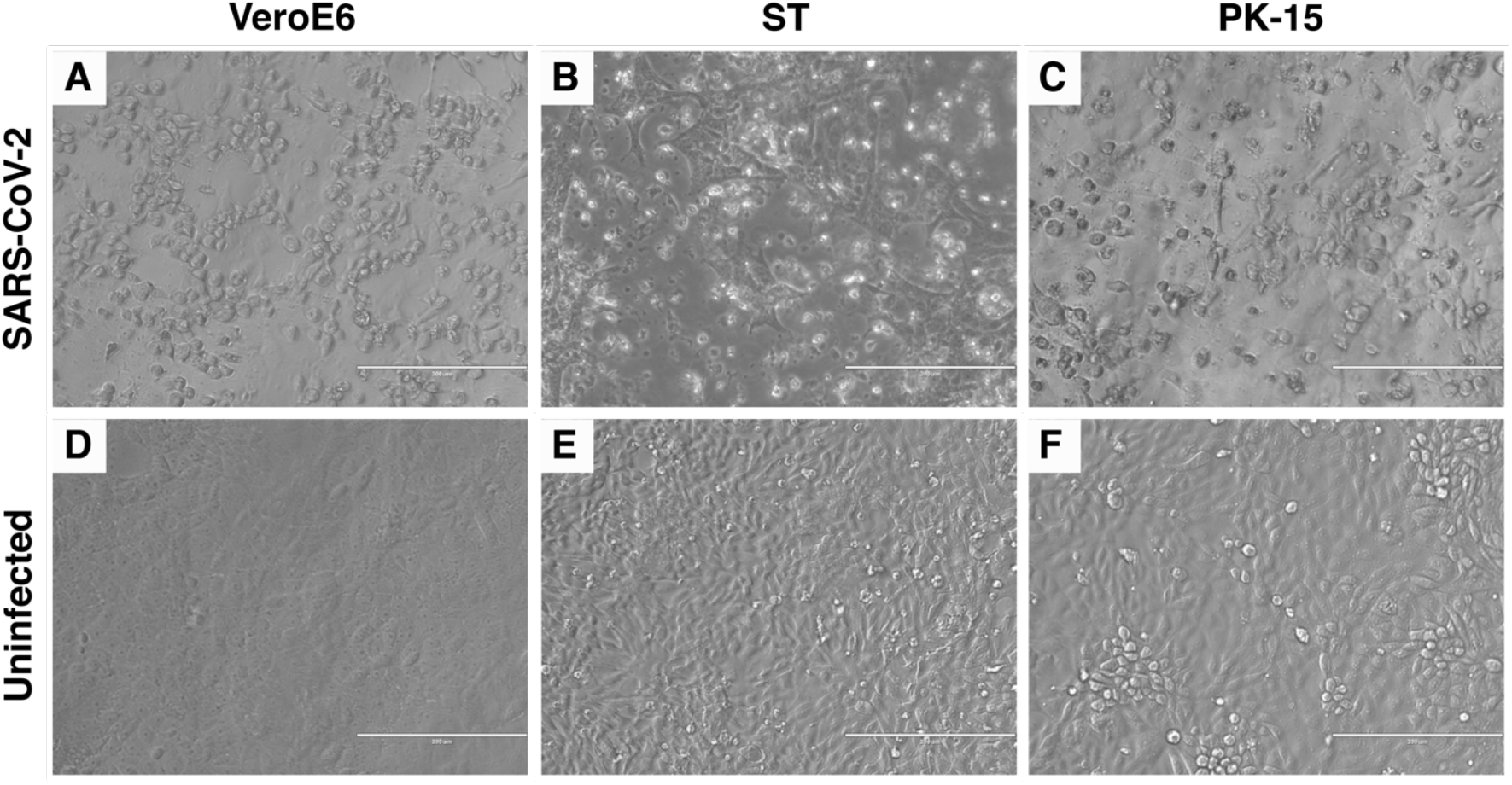
Cytopathic effect (CPE) of SARS-CoV-2 in Swine Testicle (ST) and Porcine Kidney (PK-15) cells. SARS-CoV-2 USA-WA1/2020 isolate was passaged in ST and PK-15 porcine cells. Passage two in ST cells (B) and passage four in PK-15 cells (C) resulted in clear CPE, similar to that observed in permissive VeroE6 cells (A). No CPE is observed in the uninfected VeroE6 or porcine cells lines (D, E, F).

### Oral/intranasal/intratracheal inoculation of pigs with SARS-CoV-2

To determine the effect of SARS-CoV-2 infection in domestic pigs, nine six-week-old SARS-CoV-2 seronegative piglets were inoculated with a total of 1 x 10^6^ TCID_50_ of the USA-WA1/2020 isolate, which was passaged once in swine ST cells (Figure 1). The challenge material (total 4 mL) was administered orally (1 mL), intranasally (1 mL; 0.5 mL each nostril) and intratracheally (2 mL) after sedation of the animals. At 1-day post challenge (DPC), six uninoculated sentinel contact pigs were co-mingled with the principal inoculated animals (3 animals per pen). Daily rectal temperatures were recorded for each pig and clinical signs were monitored daily, including observations for signs of lethargy, hyporexia, respiratory distress (coughing, labored breathing, nasal discharge), and digestive issues (diarrhea or vomiting). No significant temperature elevation or change in rectal temperature was observed in the principal inoculated nor sentinel contact pigs throughout the study (Figure 3). Moreover, no obvious clinical signs were observed for any of the principal inoculated nor sentinel pigs throughout the 21-day observation period.

**Figure 3.**
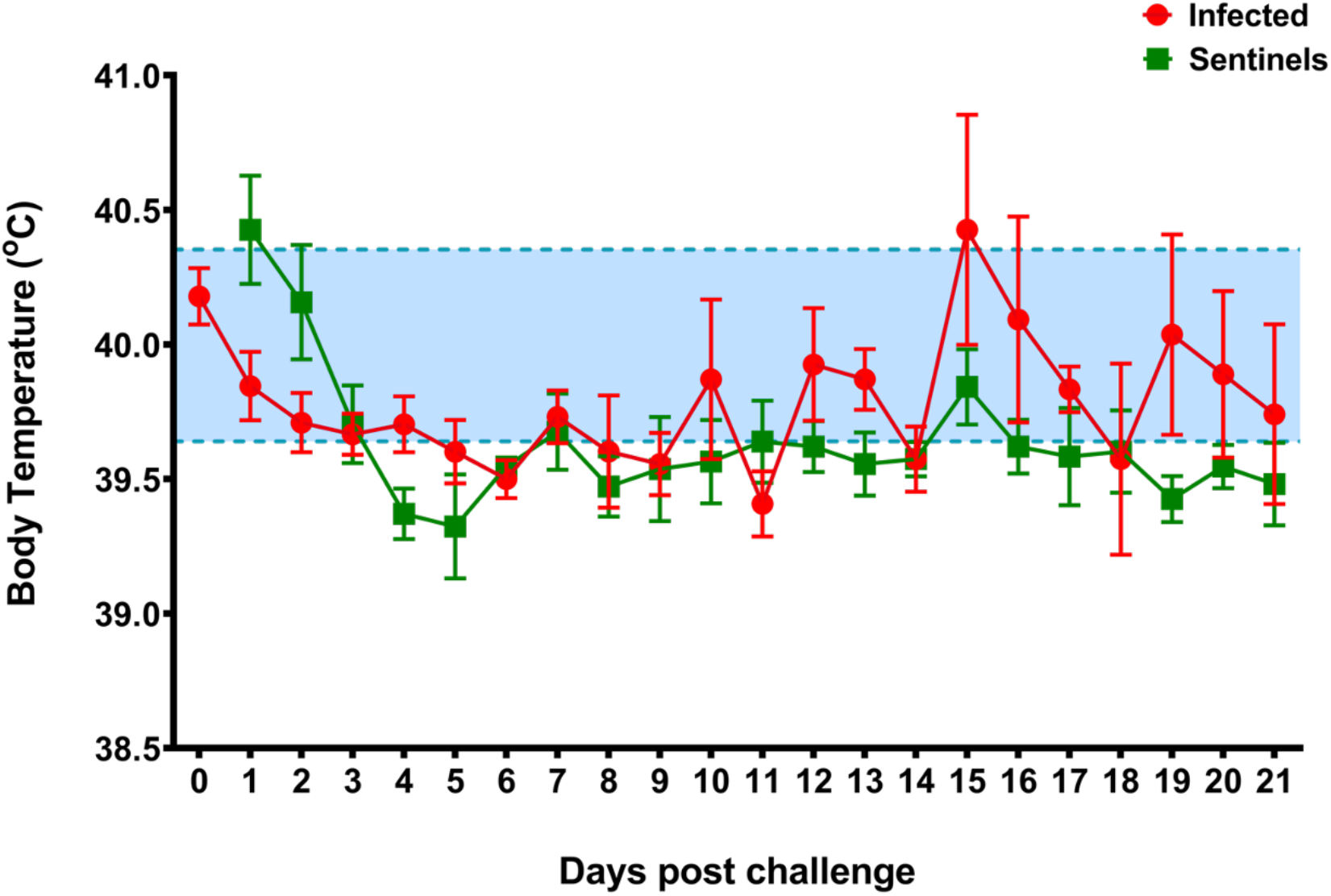
Average daily rectal temperatures of SARS-CoV-2 inoculated and sentinel pigs. Daily average rectal temperatures of pigs inoculated orally, intranasally, and intratracheally with SARS-CoV-2 (red) and co-mingled sentinel pigs (green) showed no significant change over the course of the experiment. The baseline temperature (blue; 39.6°C to 40.4°C) was determined from all pigs before infection.

To detect viral replication in the principal and sentinel pigs, clinical samples were subjected to RT-qPCR to detect the SARS-CoV-2 N gene (Table 2) [35]. Blood and oropharyngeal, nasal, and rectal swabs were collected at various time points throughout the study and upper and lower respiratory tract tissues were collected at *post-mortem* examinations on 4, 8 and 21 DPC. RT-qPCR failed to detect any viral RNA in any swab or blood sample for the duration of the study (Table 2). The only the exception was a low suspect positive result in a nasal swab at 1 DPC in a principal inoculated pig #161, for which one of two qPCR replicates yielded a low fluorescent amplification curve with a Ct of 37 (Table 2). Moreover, viral RNA was not detected in any lung sample collected at *post-mortem* examination on 4, 8 and 21 DPC (Table 2). In addition, gross and histopathological analysis of trachea and lung from the principal challenged pigs did not reveal the presence of any obvious pathological lesions (Table 3, Figure 4). These results indicate that SARS-CoV-2 failed to replicate in the respiratory and digestive tract as well as the blood in orally/intranasally/intratracheally inoculated pigs throughout an observation period of 21 days. This is confirmed by the fact that the principal infected pigs failed to transmit SARS-CoV-2 to co-mingled sentinel animals.

**Table 2.** Summary of RT-qPCR results.

| Group | Nasal Swabs | Oropharyngeal Swabs | Rectal Swabs | Blood | Lung |
| --- | --- | --- | --- | --- | --- |
| Inoculated<sup>1</sup> | -* | - | - | - | - |
| Sentinel<sup>2</sup> | - | - | - | - | - |
| Uninfected<sup>3</sup> | - | - | - | - | - |

(-) = negative
<sup>1</sup>Swabs/blood were tested from samples on 0, 3, 5, 7, 10, and 14 DPC. Lung tissue was collected 4, 8, and 21 DPC.
<sup>2</sup>Swabs/blood were tested from samples on 0, 3, 5, and 10 DPC. Lung tissue was collected on 21 DPC
<sup>3</sup>Swabs/blood were tested on 0 DPC. Lung tissue was collected on 3 DPC for these uninfected controls.
\*One pig (#161) had a ct signal of 37.72 ( $3.82 \times 10^4$ copy number/mL) for 1/2 of RT-qPCR wells on 1 DPC.

**Table 3.** Macroscopic lesions of total lung (%).

| Group | Day of Necropsy | Average Score |
| --- | --- | --- |
| Inoculated | 4 DPC | 1.6 |
|  | 8 DPC | 1.1 |
|  | 21 DPC | 2.9 |
|  | All pigs | 1.9 |
| Sentinels | 21 DPC | 3.4 |
| Uninfected | 3 DPC | 0.1 |

**Figure 4.**
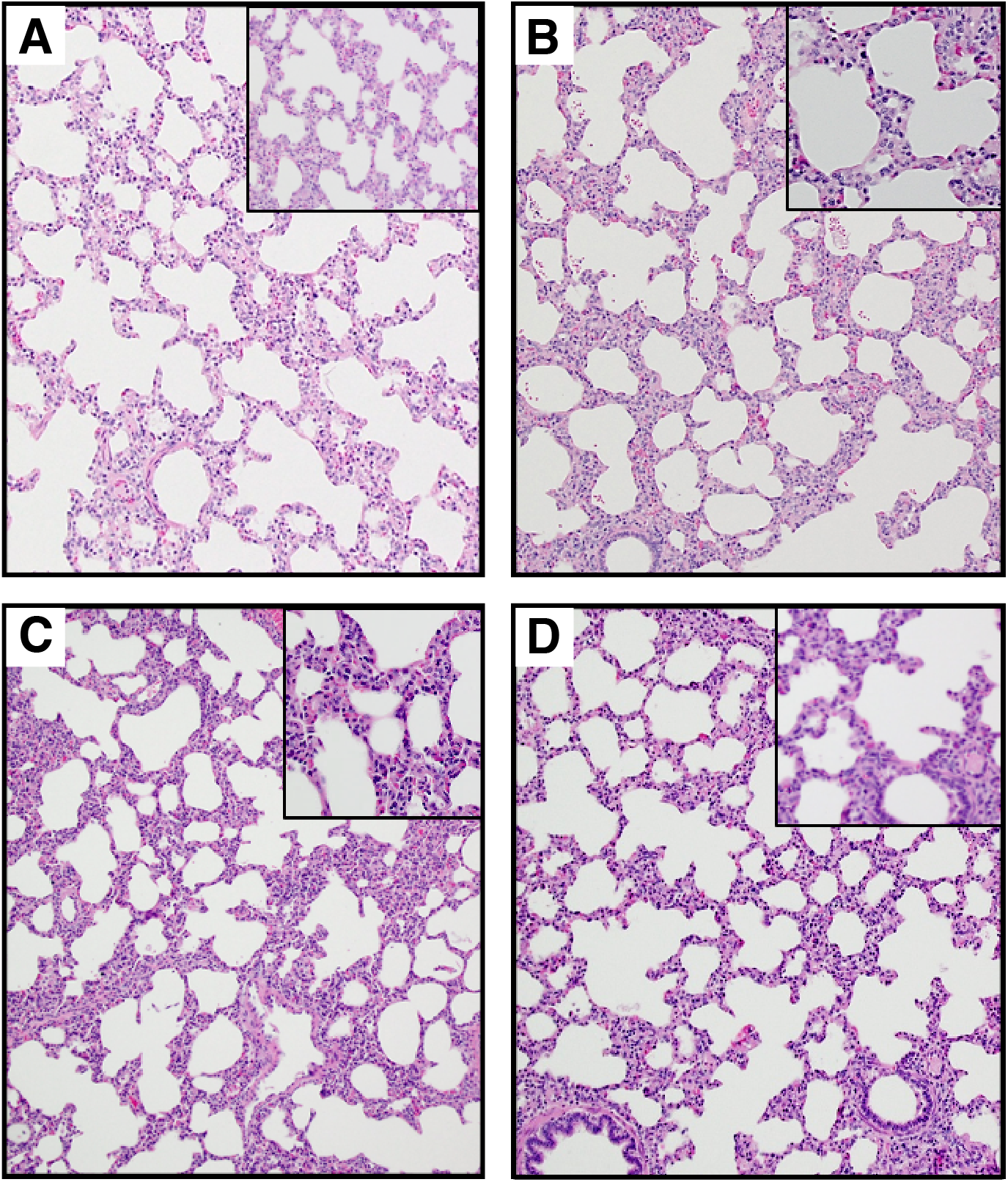
Histopathological analysis of pig lung tissue. Lung tissue sections were stained with hematoxylin and eosin for histopathological evaluation. (A) Uninfected negative control pig #210, (B) Infected pig #161 4 DPC, (C) Infected pig #803 8 DPC, (D) Infected pig #193 21 DPC. No significant histopathology was observed. Magnification is 10x for main images and 20x for inserts.

### Absence of immune response in SARS-CoV-2-inoculated pigs

To determine whether the orally/intranasally/intratracheally inoculated pigs developed an immune response against SARS-CoV-2 antigens, sera collected from infected pigs at various time points post infection was subjected to indirect ELISAs to detect IgG and IgM antibodies reactive against the receptor-binding domain (RBD) of the SARS-CoV-2 spike protein and the SARS-CoV-2 nucleocapsid (N) protein. Cumulatively, the results from these assays indicated that the infected pigs did not develop an IgG or IgM immune response against either of the SARS-CoV-2 antigens at any point throughout the study (Figure 5). The sera from one sentinel pig (#894) did show transient IgM reactivity on 3 and 5 DPC against N and RBD and an isolated IgG reactivity against RBD on 3 DPC. In addition, neutralizing antibody experiments performed with sera collected from principal inoculated and sentinel contact pigs necropsied at 14 and 21 DPC, revealed that none of the pigs developed neutralizing antibodies against SARS-CoV-2. Overall, these results indicate that SARS-CoV-2 is unable to replicate and generate an immune response in pigs upon oral/intranasal/intratracheal inoculation. In addition, transmission to sentinel contact animals was not possible.

**Figure 5.**
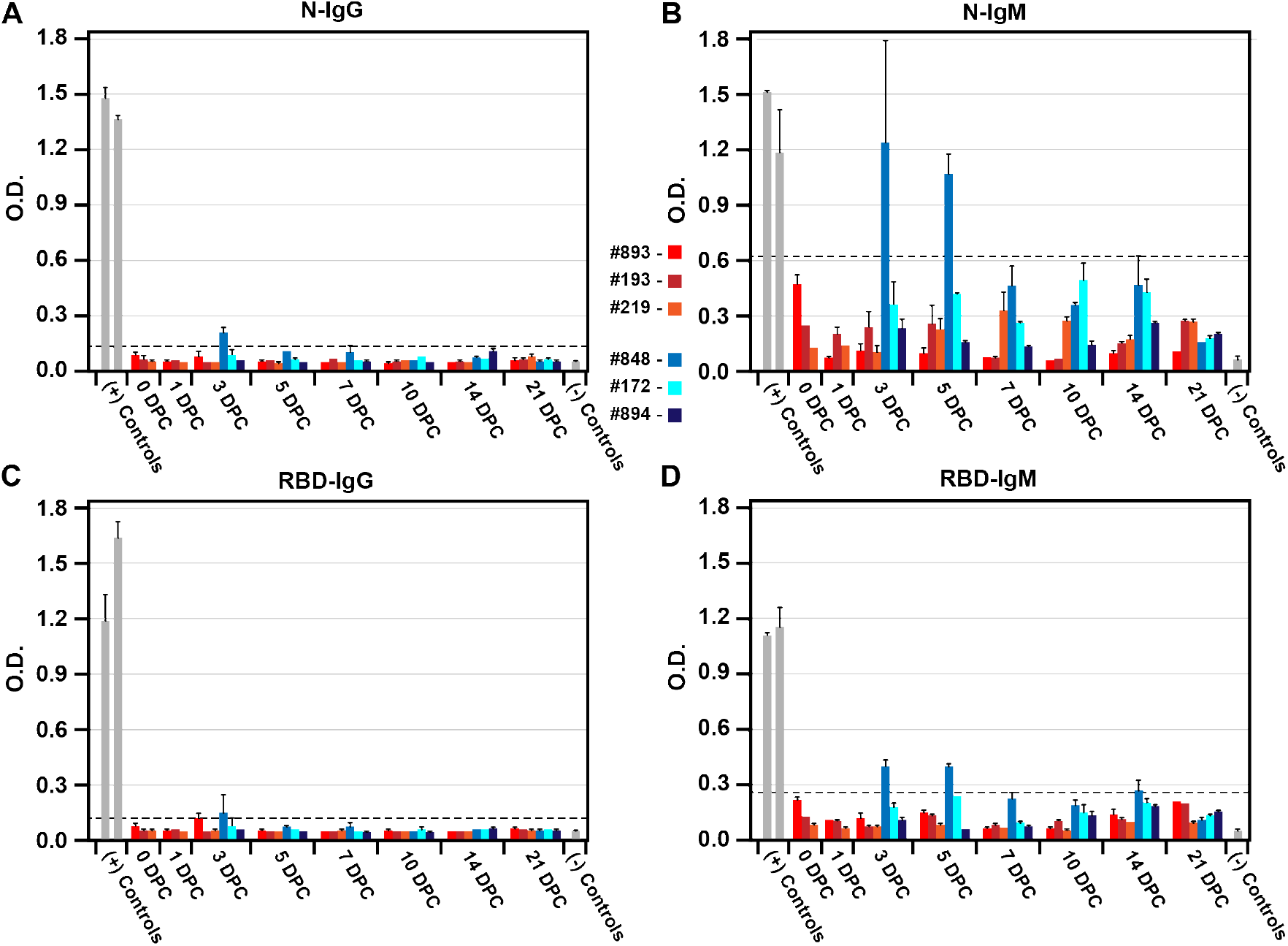
Immunological response in pigs infected orally/intranasally/intratracheally with SARS-CoV-2 and sentinel contact pigs. Indirect ELISAs were performed against the SARS-CoV-2 antigens N (nucleocapsid protein [A, B]) and RBD (Spike protein receptor binding domain [C, D]) to detect an antigen-specific IgG (A, C) or IgM (B, D) antibodies. Sera reactivity was determined for three principal infected pigs (#893, #193, #219) and three sentinel pigs (#848, #172, #894). The cutoff for a positive sample was determined by +3 standard deviations of 0 DPC samples (dotted line). Feline SARS-CoV-2-specific antibodies were used as a positive control from a separate study (left bar, [38]). Porcine positive control sera for IgG/IgM-specific antibodies were ASFV-infected pig sera detecting the ASFV-p54 antigen (right bar). Uninfected pigs were used as a negative control. O.D. – optical density.

## Discussion

SARS-CoV-2 is a zoonotic agent, and a detailed understanding of the susceptibility of various animal species to SARS-CoV-2 is central to controlling its spread [16,17]. In addition, the development of animal models that emulate COVID-19 in humans is essential for pre-clinical testing of novel vaccines and therapeutics [20]. In this study, we inoculated nine pigs with a high dose of SARS-CoV-2 that was passaged once in porcine cells. Simultaneous oral/intranasal/intratracheal inoculation did not result in any detectable viral RNA in the blood, the oral/nasal/rectal cavities, or the lungs. Also, none of the co-mingled, sentinel contact pigs shed viral RNA. Moreover, a virus-specific immune response characteristic of infection was not observed within the 21-day study period in the principal infected or sentinel pigs. The transient nature of the IgM and IgG response observed in pig #848 could indicate cross-reactivity of antibodies directed against a porcine coronavirus such as porcine epidemic diarrhea virus [40]. Such antibodies could be maternally derived and therefore transient as the lack of SARS-CoV-2 specific reactivity by the end of the study might suggest. In contrast to previous SARS-CoV-2 swine studies [22,26], the present study used a more stringent inoculation procedure (intratracheal and oral, in addition to intranasal) and 1 log higher titer of virus inoculum (10^6^ vs 10^5^). In addition, the inoculum in the present study was passaged once in porcine ST cells. These results, combined with previous intranasal pig inoculation studies [22,26], indicate that pigs seem to be resistant to SARS-CoV-2 infection, are unlikely to be a SARS-CoV-2 carrier animal species, and are also not suitable as an animal model for research.

The results of the present and previous SARS-CoV-2 inoculation studies in pigs are intriguing in light of the findings that the porcine ACE2 receptor seems highly compatible with the SARS-CoV-2 RBD, suggesting that pigs could be susceptible to SARS-CoV-2 infection [2,31]. Pigs are susceptible to both experimental and natural infection with SARS-CoV-1 [29,30]. However, the experimental SARS-CoV-1 infection was via simultaneous intranasal/oral/intraocular/intravenous inoculation [29], thus the actual route(s) of SARS-CoV infection cannot be determined. Recently, several porcine cell lines have been shown to be permissive to SARS-CoV-2 infection [26,33]; in addition, single-cell screening studies showed that porcine ACE2/TMPRSS2 expression are compatible with infection [32]. In contrast to previous reports that some porcine cell lines are susceptible to SARS-CoV-2 infection, but show no CPE [26,33], we found that both ST and PK-15 cell lines are susceptible to infection and observed CPE after two or four passages, respectively. The absence of SARS-CoV-2 replication and transmission in the present and two previous pigs studies [22,26] seems to lessen the need to monitor pig populations for SARS-CoV-2 during the ongoing pandemic.

However, the evidence described above suggests pig susceptibility should not be disregarded, because all pig studies to date have used rather young pigs and commercially available pig breeds/genetics. We also have to be aware that unforeseen genetic changes in the SARS-CoV-2 genome may result in a better compatibility of the virus for pigs in the future.

Pigs are considered to be an excellent model for studying human infectious diseases based on their relatedness to humans in terms of anatomy and immune responses and they have been found to be much more predictive for the efficacy of therapeutics when compared to rodent models [41]. However, the results presented here indicate that pigs are not a suitable preclinical model for SARS-CoV-2 pathogenesis studies and the development and efficacy testing of therapeutics and/or vaccines. A recently available article indicates that while pigs are not susceptible to SARS-CoV-2 infection, neutralizing antibody responses were detected in pigs infected via intramuscular or intravenous inoculation routes [42]; this indicates that pigs could be used for immunogenicity studies related to SARS-CoV-2. However, the use of pigs to monitor SARS-CoV-2 immune responses must be careful to screen for cross-reactive maternal antibodies derived from other coronaviruses [43]. Alternate pre-clinical animal models, namely non-human primates, Syrian hamsters, transgenic or transduced mice expressing human ACE2, ferrets, or even cats need to be considered to gain additional insights into SARS-CoV-2 pathogenesis and virulence. Comprehensive characterization of SARS-CoV-2 pathogenesis in preclinical animal models and the establishment of standardized infection and testing protocols will be crucial for the development of much-need countermeasures to combat COVID-19.

## Acknowledgements

We gratefully thank the staff of KSU Biosecurity Research Institute, the histological laboratory at the Kansas State Veterinary Diagnostic Laboratory (KSVDL), the CMG staff and Gleyder Roman-Sosa, Yonghai Li, Konner Cool, Emily Gilbert-Esparza, and Chester McDowell at KSU. The following reagent was obtained through BEI Resources, National Institute of Allergy and Infectious Diseases (NIAID), National Institutes of Health (NIH): SARS-CoV-2 Virus strain USA-WA1/2020 (catalogue # NR-52281).

